# An Open-source, Cost-Efficient Excitation Module for Single-molecule Microscopy

**DOI:** 10.1101/2022.02.28.482236

**Authors:** Dylan R. George, Benjamin Ambrose, Ashley J. Cadby, Timothy D. Craggs

## Abstract

In fluorescence microscopy, scientific-grade laser diodes are key components and contribute a considerable expense to the total cost of the microscope setup. Existing open-source laser excitation modules are mainly designed for super-resolution imaging and have not been fully characterised to meet the requirements of single-molecule spectroscopy measurements. In this paper we introduce an open-source, cost-efficient excitation module that can be used for confocal, accurate single-molecule FRET, fluorescence correlation spectroscopy (FCS) and low-power super-resolution techniques. The module delivers two diode lasers (520 nm and 638 nm) via a single-mode fibre, with stable power output (<1% deviation), which can be modulated on the microsecond timescale. Here, we benchmark these lasers against smFRET standards, and recover consistent diffusion coefficients from FCS measurements, thereby demonstrating their suitability for a range of single-molecule spectroscopic experiments, whilst reducing the overall costs of our open-source smFRET instrument (the smfBox) by ~50%, making these types of experiments even more accessible to the widest possible userbase.

## 1. Introduction

Fluorescence microscopy encompasses a range of biophysical imaging techniques such as epi-fluorescence, confocal and total internal reflection fluorescence (TIRF) microscopy. Fluorescent samples are excited by a high energy light source at specific wavelength (typically a laser) and dichroic filters are used to separate the excitation light from the fluorescence emission to provide higher contrast than in conventional microscopy. Recently, some labs have provided a gateway into this area for non-specialists by publishing open-source microscopy platforms and software packages.^[1]^ A range of fully open-source microscopy platforms are now currently available including: the miCube, capable of super-resolution microscopy, TIRF and LED brightfield microscopy;^[2]^ the openSPIM, a light-sheet microscope; and the LifeHack microscope, which provides single-molecule localisation microscopy (SMLM) and live cell imaging.^[3]^ Open-source projects also include common microscope components such as microscope fluidic systems, lasers and stages.^[4–7]^ Open-source software for super-resolution imaging has also been developed.^[8–10]^ Previously, we introduced the smfBox, an open source single-molecule FRET microscopy platform to combat the high instrument costs and a lack of open-source hardware that has limited this technique’s broad application by non-specialists.^[11]^ In general, the lasers for fluorescence microscopy applications incur a considerable expense. Since multiple fluorophores with different spectra are often used to differentiate biological components, multiple lasers are employed to excite them. However, these laser setups, whether bought commercially or custom-made, often make up a large proportion of the overall cost of modern microscopes. Many fluorescence applications require different excitation specifications (in terms of laser powers and pulsed / alternation time periods) increasing the number of lasers needed. This makes it especially difficult to provide a universal excitation module that is suited to all microscopy applications. Table 1 provides a short list of some of the most common fluorescence applications, and their excitation requirements for normal operation.

**Table 1.** Common fluorescent applications, their excitation system requirements and primary expense.^[12–15]^

| Technique | Min. Number of Lasers | Approximate Power Range | Pulsed Time Period | Key Requirements |
| --- | --- | --- | --- | --- |
| **Confocal Single-molecule FRET with Alternating Laser Excitation (ALEX)** | 2 | (Hundreds of) $\mu\text{W}$'s | $\mu\text{s}$ | Multiple Lasers |
| **TIRF Single-molecule FRET with Alternating Laser Excitation (ALEX)** | 2 | (Hundreds of) $\mu\text{W}$'s | $\mu\text{s}$ | Multiple Lasers |
| **Fluorescence correlation spectroscopy (FCS)** | 1 | (Hundreds of) $\mu\text{W}$- (Tens of) $\text{mW}$ | CW | Stable power/ Fast modulation |
| **Nano-second FCS (nsFCS)** | 1 | (Hundreds of) $\mu\text{W}$- (Tens of) $\text{mW}$ | CW | Stable power/ Fast modulation |
| **Fluorescence cross-correlation spectroscopy (FCCS)** | 2 | (Hundreds of) $\mu\text{W}$- (Tens of) $\text{mW}$ | CW | Multiple Lasers |
| **Single-molecule localization microscopy (SMLM)** | 1 | (Ten's of) $\text{mW}$'s - (Hundreds of) $\text{mW}$ | CW | High laser power |
| **Stimulated emission depletion (STED) microscopy** | 2 | (Hundreds of) $\text{mW}$'s | ps-fs | High laser power, multiple lasers |
| **Light-sheet microscopy (LSM)** | 1 | $\mu\text{W}$'s | CW | Low laser power |
| **Coherent raman scattering (CARS)** | 2 | $\text{mW}$'s | ps | Multiple lasers |
| **Total internal reflection fluorescence (TIRF)** | 1 | (Tens of) $\text{mW}$'s | CW | Low laser power |
| **Confocal** | 1 | $\mu\text{W}$'s-W's | CW | Low laser power |
| **Structured illumination microscopy (SIM)** | 1 | $\mu\text{W}$'s-W's | CW | Low laser power |
| **Widefield** | 1 | $\mu\text{W}$'s-W's | CW | Low laser power |

Over the last decade, a few open source, affordable laser systems have been developed (Table 2). However, these have mainly been targeted towards super-resolution imaging; For example, the low-cost, continuous wave Laser Engine (from Schroder et al.) which provides a homogeneous, scatter-free laser beam through a multi-mode fibre vital for single-molecule localisation microscopy.^[5]^ The NicoLase (from Nico et al.) caters to a broader range of applications with many different lasers in the visible spectrum and provides a full characterisation of the alternation of the lasers required for many single-molecule applications, albeit using higher-end laser diodes which may significantly add to the cost of the setup.^[6]^

**Table 2.** A list of current open-source laser excitation modules; their properties and their primary application.^[5–6]^

| Author | No. of Lasers | Wavelengths | Power | Modulation | Fibre | Key Application |
| --- | --- | --- | --- | --- | --- | --- |
| **Schroder *et al.*** | 3 | 405 nm, 488 nm, 638 nm (561 nm option) | (Hundreds of) mW | N/A | Multi-mode | Single-molecule localisation microscopy |
| **Nico *et al.*** | 5 | 405 nm, 490 nm, 517 nm, 561 nm, 640 nm | (Hundreds of) mW's | Arduino Uno | Single-mode & Multi-mode | Fluorescence/super-resolution |
| **This Work** | 2 | 520 nm, 638 nm | (Tens of) mW's | NI DAQ board | Single-mode | Single-molecule FRET, FCS |

While these open-source excitation modules are available, many researchers have not incorporated them into single-molecule spectroscopy methods as a full characterisation of their properties and suitability for this type of application has been lacking. This is particularly true for single-molecule FRET applications, in which precise modulation on the microsecond timescale and stable laser powers are crucial to achieve accurate FRET measurements. Here, we present, an open source, affordable and fully characterised excitation module capable of accurate single-molecule FRET and fluorescence correlation microscopy. This excitation module overcomes the cost of commercial pulsed laser systems whilst still offering high laser stability and precise modulation. This module reduces the cost of the smfBox by ~50%, allowing more researchers to harness the power of single-molecule FRET in their research, whilst simultaneously reducing the cost and time taken to build an excitation system.

In single-molecule FRET, alternating laser excitation (ALEX) provides information about the presence of both donor and acceptor fluorophores.^[16]^ In confocal smFRET, molecules freely diffuse through a near-diffraction limited spot formed by focussing expanded, collimated laser light through an objective lens. During the transit of this confocal volume, donor and acceptor fluorophores attached to an individual molecule are alternately excited by two lasers, leading to the emission of a burst of photons. These bursts can then be classified by their FRET efficiency and stoichiometry, an additional dimension, which defines the presence of the acceptor fluorophore (see methods). By using ALEX, donor only, acceptor only and dual labelled populations can be identified, and all the correction factors determined for generating accurate FRET efficiency values (see methods).^[17, 18]^ Here we provide a full excitation module characterisation and explain in depth how to alternate these laser modules to achieve accurate single-molecule FRET (ALEX) and fluorescence correlation spectroscopy (FCS) measurements.

## 2. Hardware/ Methods

For single-molecule FRET (ALEX) the laser diodes need to be modulated at a frequency of 20 kHz at the wavelengths of 515 nm and 638 nm. Additionally, we require laser powers of around a few hundred microwatts at the objective lens. The system also needs to be coupled to a single-mode fibre to provide a single excitation spot. The single-mode fibre provides a Gaussian beam profile for forming a high-quality confocal excitation volume.

The chosen lasers (520 nm at 100 mW and 638 nm at 700 mW, Lasertack) met all these requirements and were chosen primarily for their low cost (< € 500). These wavelengths are popular for single-molecule FRET setups as they efficiently excite a range of commonly used donor (520 nm) and acceptor (638 nm) fluorophores. Although the power of the lasers is an order of magnitude higher than the requirements for single-molecule experiments, a single-mode fibre as previously mentioned was required and so we anticipated a low coupling efficiency. However even with the achieved coupling efficiency of around 4 % (638 nm laser) and 25 % (515 nm laser) a power of 26 mW and 25 mW was delivered out of the fibre, higher than the requisite power.

The laser modules were mounted on aluminium heatsinks (100 mm x 100 mm x 30 mm, Fischer Electronik). These heatsinks were then mounted onto an 300 mm x 300 mm optical breadboard (MB3030/M, Thorlabs). The laser modules were aligned using two kinematic mirrors (KM05/M, Thorlabs) which steered the beam onto a dichroic mirror (DMLP567T, Thorlabs). The dichroic mirror combined the laser beams before a Fibreport (PAF2A-A15A, Thorlabs) coupled the beam into a single-mode fibre (P3-488PM-FC-10, Thorlabs).

To control the lasers, a modulation voltage of 5 V was delivered to the lasers at a frequency of 20 KHz. A National Instruments (NI) DAQ board was used to deliver the modulating voltage signal, controlled using the smOTTER software (available on GitHub, https://github.com/craggslab/smfBox). Alternatively, a simple LABVIEW program (supplied in the supplementary material) can be used.

## 3. Results and Discussion

### 3.1. Power and power stability

We conducted a full characterisation of the laser modules to check whether it was able to perform single-molecule experiments. To check whether the module needed excitation filters the wavelength spectrum of both the 520 nm and 638 nm laser modules was recorded (Figure 2). The wavelength of the green laser diode ranged from 512 nm – 526 nm and the spectrum of the red laser diode 630 nm – 644 nm (Figure 2). The resulting spectrum from both lasers was within the range expected and was sufficiently tight for single-molecule experiments without any excitation filters.

**Figure 1.**
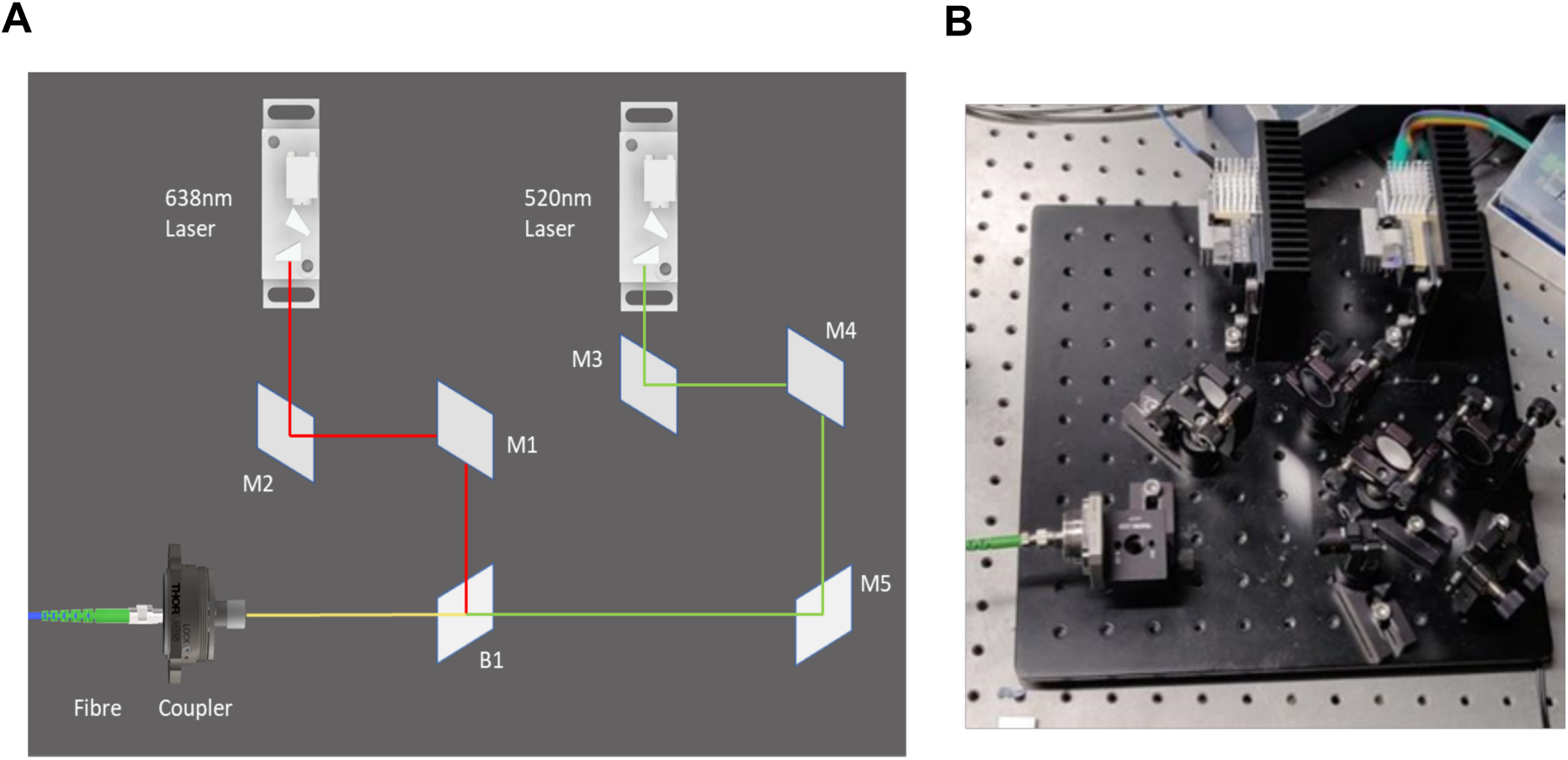
A) A schematic showing the excitation pathway of the module. Red and green line shows the path of the laser beams travelling towards the coupler. The mirrors (M1-M5) are used to align the laser beam. A dichroic beamsplitter (B1) reflects the red laser beam and allows the green laser beam to travel through. Both laser beams are then coupled into a fibre via a coupler. Further details (including part numbers) are found in supplementary information 1.1) B) A photograph of the finished excitation module showing the excitation in the same orientation as the schematic.

**Figure 2.**
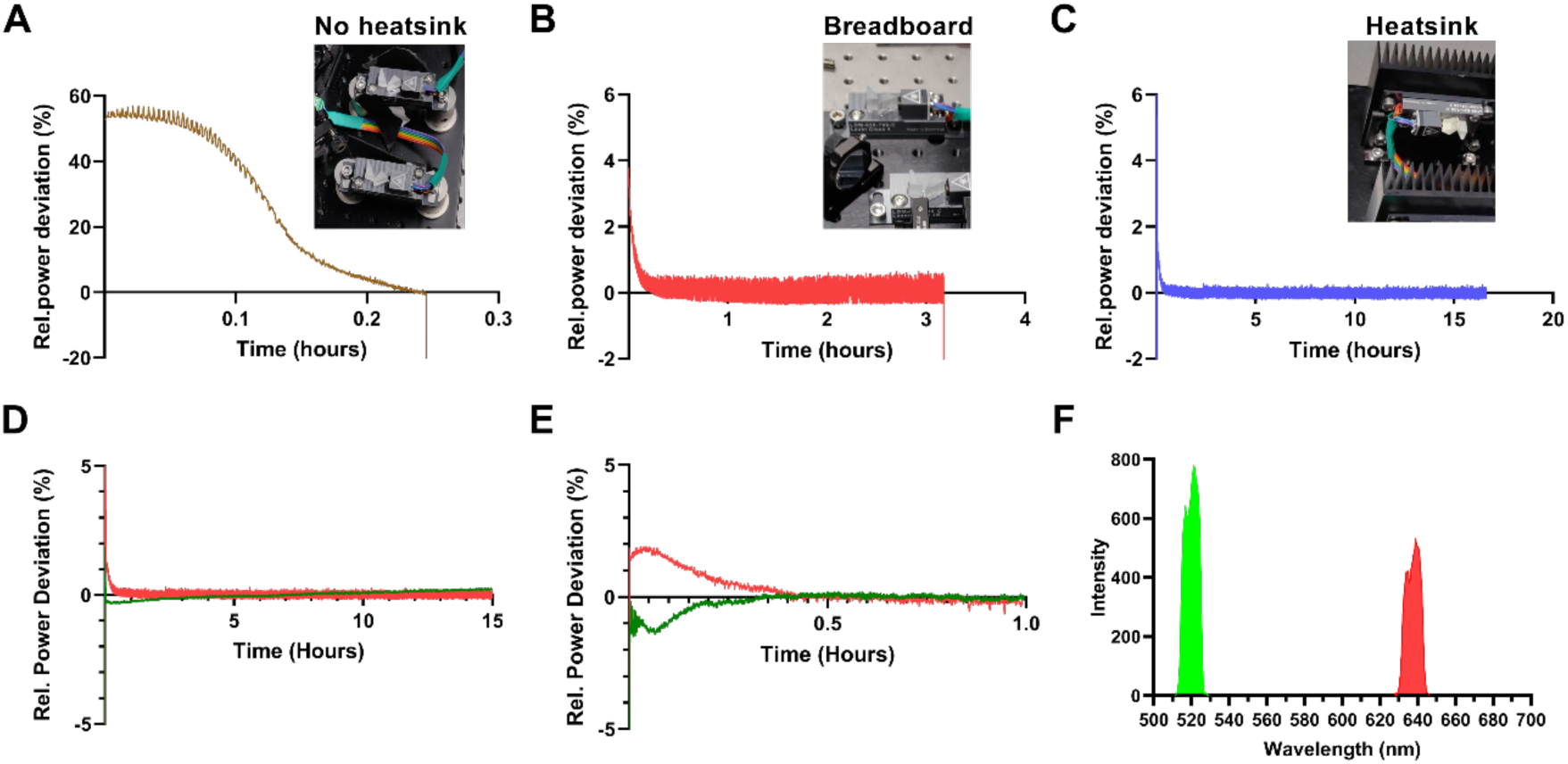
A-C) The power stability of the lasers with different heat dissipation mountings: posts alone (A), breadboard (B) and heatsinks (C) D) Relative power stability for 520 nm (green line) and 638 nm (red line) lasers from switch on, under continuous wave mode E) Relative power stability for both lasers from switch on, under 20 kHz alternation F) The wavelength spectrum of the 520 nm (green) and the 638 nm (red) lasers.

Since single-molecule measurements are taken on short timescales (ms), inconsistencies in laser power on this timescale can give rise to differences in the amount of fluorescent emission detected. This is particularly important for FCS, where instabilities in laser power can affect the accuracy of results, but also needed for smFRET to obtain good quality data. In the initial design, the laser modules were placed on pedestal posts. A power stability assessment of the red laser module on these posts revealed that the laser power was particularly unstable. Additionally, the laser module switched off after less than 30 mins, below the acquisition period required for many single-molecule FRET experiments. The laser modules were then mounted on an aluminium breadboard to improve heat dissipation. On the breadboard, the laser module had a mean relative stability percentage of 0.083 % with a standard deviation of 0.20 % and switched off after just over 3 hours. Finally, the laser modules were mounted on heatsinks. On the heatsinks, the laser module had a mean relative stability percentage (M) of −0.03 % with a standard deviation (SD) of 0.10%. Furthermore, the laser beam remained stable and active for over 15 hours, well above the acquisition time required for most single-molecule FRET experiments. The power stability test was repeated for the green laser (mounted on the heat sink) which showed a similar performance (M= 0.004 %, SD=0.14 %, Figure 2E). The laser modules were also alternated on and off for time periods of 45us on and 55us off using the single-molecule FRET acquisition software (smOTTER). After a 30 minute warm up time the laser modules showed high stability (Red - M = −0.08%, SD = 0.11 %, Green - M = 0.002 %, SD = 0.07 %), sufficient for single-molecule FRET measurements.

### 3.2. Alternation rise and fall times for ALEX

To achieve accurate FRET values, alternating the red and green lasers at a frequency of 20 kHz was required. The lasers were alternated by the single-molecule FRET acquisition software (smOTTER). The output of the fibre was coupled to a photodiode to measure the laser power with high (ns) time resolution. Initial results showed that the lasers in the excitation module could indeed be successfully modulated at the required microsecond timescales (Figure 3A). We next investigated the rise and fall times of the lasers and compared them to the commercially available laser combining system currently used on the smfBox. The rise times for the excitation module were recorded as 8 μs for the 520 nm laser and 6 μs for the 638 nm laser (defined as the time taken to reach the plateau in signal). Both red and green lasers in the excitation module had a recorded fall time of 5.5 μs. For the commercially available lasers, these times were slightly shorter with a recorded rise time of 4 μs for the green laser and 4 μs for the red laser. Similarly, the fall times were recorded as 4.5 μs for the green laser and 3.5 μs for the red laser. While the rise and fall time of the excitation module performed slightly worse than the currently used lasers on the smfBox, crucially, these values were still within the acceptable range for obtaining high quality single-molecule FRET data.

**Figure 3.**
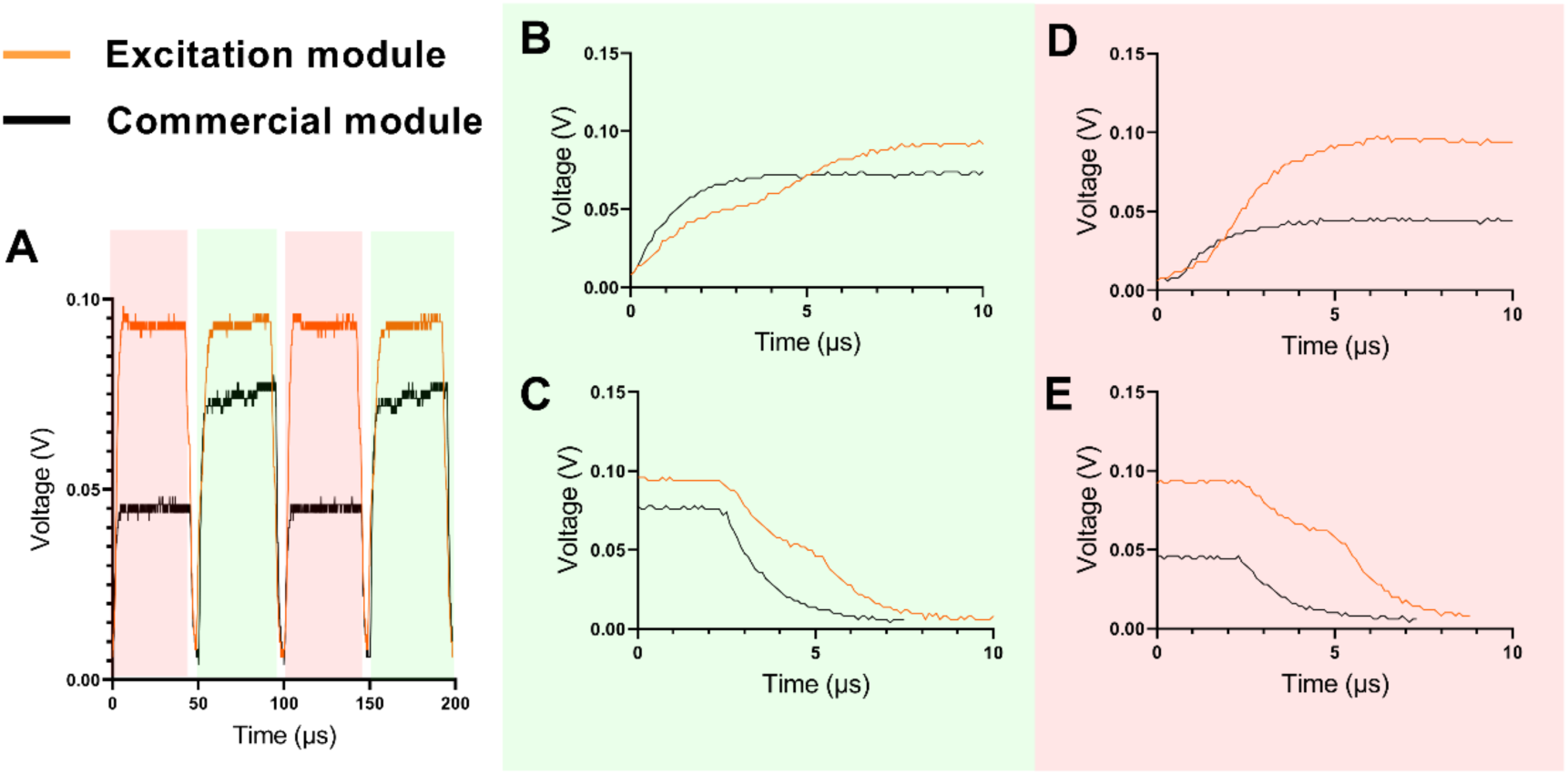
A) Time-resolved measurements of the laser power measured by a photodiode B-F) Time trace of a commercially available laser alternation (black) and the excitation setup (orange) B) Green laser rise time C) Green laser fall time D) Red laser rise time E) Red laser fall time.

### 3.3. Applications

#### 3.3.1. smFRET

Since the results of the characterisation of the laser modules were sufficient for single-molecule FRET measurements, the excitation module was used on the smfBox to take smFRET measurements. Three double-stranded DNA samples were used given their well characterized FRET values in the literature.^[18]^ These samples are labelled with ATTO 550 (donor) and ATTO 647N (acceptor) with the donor dye positioned at 23bp, 15bp and 11bp away from the acceptor dye. These three DNA standards have reported FRET Efficiency values of 0.15±0.02, 0.56±0.03 and 0.76±0.015, respectively, as measured in a recent blind multi-lab study. ^[11, 18]^

The procedure for determining accurate FRET efficiencies for absolute distance measurement is well documented in the literature. To yield accurate FRET efficiency values, both the uncorrected stoichiometry and the uncorrected FRET efficiency (proximity ratio) must be determined using alternation excitation (ALEX). The uncorrected stoichiometry provides a pathway for the identification of acceptor only and donor only populations. Both the stoichiometry and uncorrected FRET efficiency are calculated via Equation 1 and 2 where D_ex_ D_em_ represent donor excitation and donor emission. Conversely, the symbols A_ex_ and A_em_ represent the acceptor excitation and acceptor emission. For instance, D_ex_A_em_ is the acceptor emission when under donor excitation.

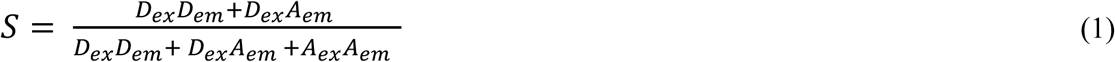

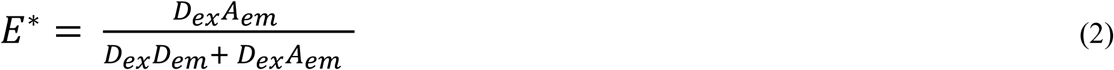

To determine these parameters, Jupyter notebooks from the smfBox GitHub along with FRETBursts were used.^[19]^ The correction factors used to find accurate FRET efficiency values were then determined. The first two correction factors α and δ were determined from the donor fluorescence leakage into the acceptor channel and the direct excitation of the acceptor with the donor laser, respectively. Using the average values for both α and δ for all three samples, the correction factors γ and β were determined. The correction factor γ takes into account the different detection efficiencies of the donor and acceptor emission photons, while β Describes their different excitation efficiencies by the system (see supplementary information).

To compare the results, Table 3 shows the FRET efficiency values of a benchmark study measuring these duplex DNA strands from 20 independent research groups (Hellenkamp et al., 2018). The table also contains the FRET values of these samples on the smfBox obtained using commercial lasers (Ambrose et al).^[11]^ By implementing the excitation setup on the smfBox FRET efficiency values for the samples 1a, 1b and 1c were 0.198±0.02, 0.599±0.17 and 0.793±0.09, respectively (Figure 4). The results were consistent with the literature values and with comparable precision to the literature. Overall, these results indicate that the excitation module can reproduce accurate FRET values.

**Table 3.**
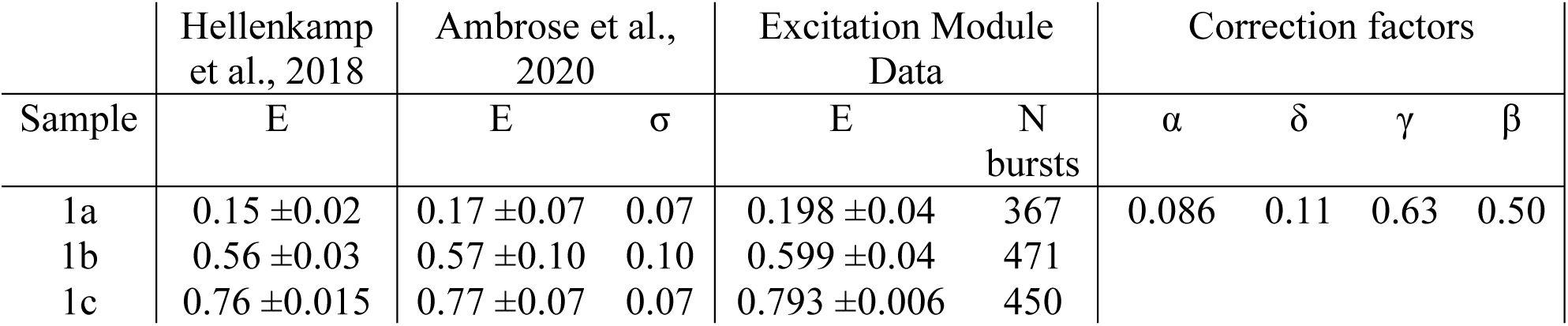
Comparison of the excitation module to literature values.

**Figure 4.**
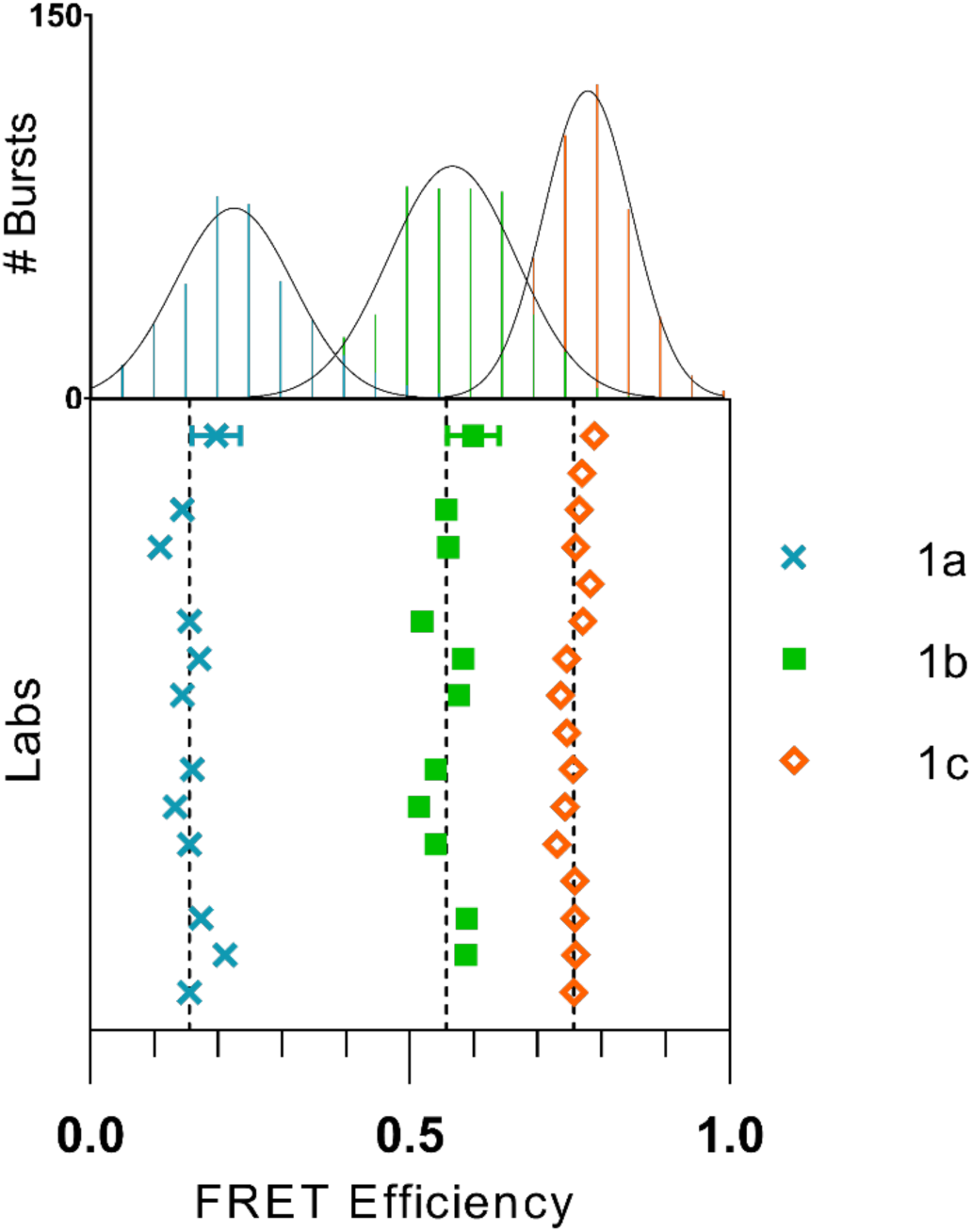
A graph showing the FRET efficiencies using the excitation module compared to the benchmark study.^[18]^ The samples 1a, 1b and 1c represent low, medium and high FRET samples, respectively. The results from the excitation layer and the FRET histogram of the excitation module are also shown at the top of the figure. These data points have error bars calculated from the standard deviation of the repeats. The 1c sample has error bars that are too small to be seen in this graph. Compared with this is the data from different labs represented by the colours in the legend. The dashed line shows the mean FRET efficiency of the labs in the benchmark study.

#### 3.3.2. FCS

Fluorescence correlation spectroscopy provides insights into the diffusion behaviour and absolute concentration of molecules by recording the fluctuation of fluorescence intensity in the observation volume. We previously measured the diffusion coefficient and molecular brightness for a duplex DNA sample on the smfBox using commercial lasers.^[11]^ To check if the excitation module could reproduce similar results the same DNA duplex sample (1a) was placed on the smfBox. Figure 5 shows the autocorrelation curve of a Rhodamine 6G sample which was used to determine the confocal volume and the diffusion coefficient of the duplex DNA sample.

**Figure 5.**
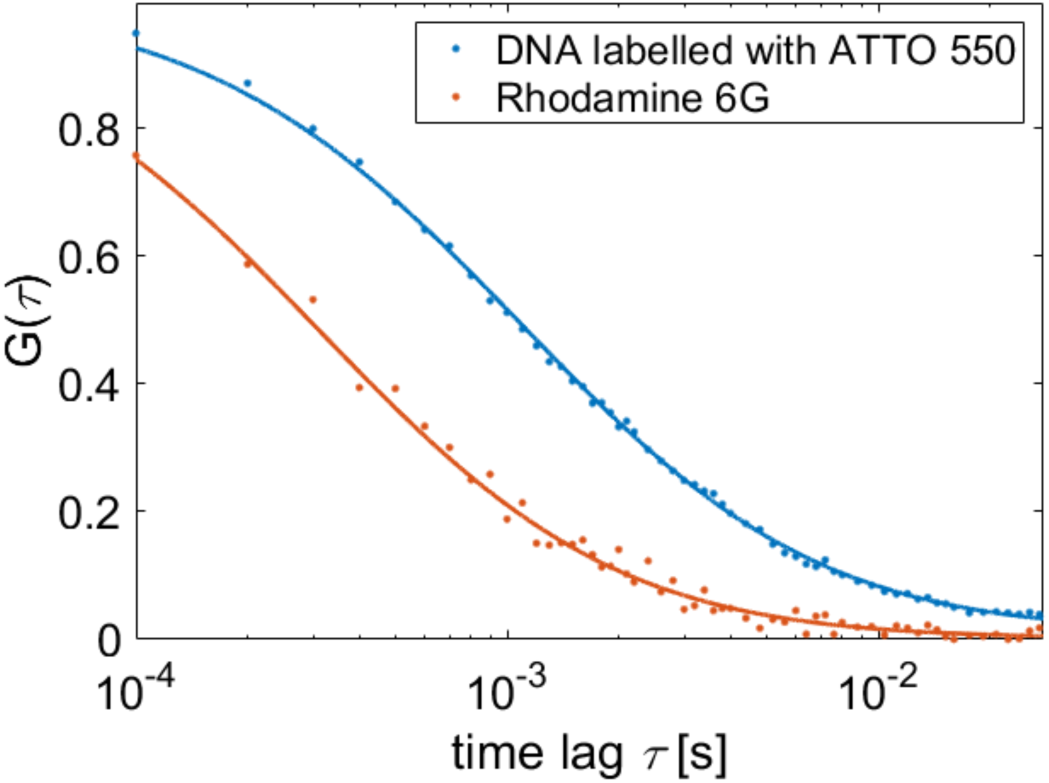
Fluorescence correlation spectroscopy measurements taken on Rhodamine 6G (orange) and a concentrated sample of ATTO 550 labelled DNA (blue) showing an autocorrelation curve.

The diffusion coefficient of the duplex DNA was measured as 118 μm^2^/s. This compared well to the previous measurement on the smfBox, using the commercial lasers, of 88 μm^2^/s.

## 4. Discussion

Here we present an affordable excitation module (costing > £ 2500) capable of fluorescence applications such single-molecule FRET and fluorescence correlation spectroscopy. This excitation module differs from existing modules, such as Schroder *et al*. and Nico *et al*. set out in the introduction, by using low-cost laser diodes inside a setup specifically designed for single–molecule spectroscopy measurements. Our data obtained using this excitation module shows that accurate FRET efficiencies and diffusion coefficients can be measured successfully. However, unlike the Nicolase and the Laser Engine, which can perform super-resolution imaging techniques, the power output of this module (~20 mW) likely falls short of the more power-hungry super-resolution applications such as STED and single-molecule localization microscopy. Due to the power and alternation requirements of microscopy applications described in the introduction we do anticipate the system will be used for FCCS, light-sheet, TIRF, confocal, SIM and widefield microscopy.

For these lasers to work effectively over the timescales and stability needed for smFRET, a series of experiments stabilising the temperature and allowing faster heat dissipation was key. Here, we were successful by mounting these lasers on heatsinks, one issue we faced was the time the lasers needed to warm-up (~20 minutes), which still arose on the second time they were switched on in close succession. However, characterising these lasers in terms of the rise times, fall times and spectrum demonstrated they could function successfully for single-molecule measurements without further modifications such as excitation filters. A simple improvement could be made by implementing a variable neutral density filter in front of each laser. This will allow the power of the lasers to be controlled.

Since using a single-mode fibre to provide a Gaussian beam is required for confocal single-molecule measurements, a drawback of using this module was the difficulty of aligning both lasers for obtaining high coupling efficiency. Care is also needed to avoid disturbing the alignment due to the sensitivity of the setup. Although, we believe that this could be improved by ensuring the optics are well secured to the breadboard and stored in a solid container. This module also makes use of an expensive NI-DAQ board to drive the microsecond alternation of the lasers. In the future, it would be more economical to implement the software to run from an Arduino or alternative. However, whilst such a solution may provide sufficiently fast electronics for alternating the lasers, the single-photon counting electronics (time-stamping photon arrival times to 10 ns bins) currently remains beyond such devices.

To conclude, this excitation module can be used for accurate single-molecule techniques at a very low-cost. This system performs well for FCS and smFRET experiments providing sufficient power and fast modulation of the lasers. We anticipate this simple to build, cost effective setup will be utilised in future single-molecule FRET and FCS setups, given the significant cost reduction (almost halving the total cost of the smfBox).

## 5. Methods

### 5.1. Characterisation of the Laser diodes

#### Characterisation of the wavelength spectrum

The 520 nm and 638 nm lasers (Lasertack) on the setup were powered on and the fibre was connected to a lens tube (SM1S20, Thorlabs). A spectrometer (Ocean Optics) was then connected to the opposite side of the lens tube facing the fibre. A neutral density filter was placed in the lens tube to avoid saturation of the spectrometer. The resultant wavelength of each of the lasers was recorded separately.

#### Characterisation of the power stability

An optical power meter was placed in a lens tube (Thorlabs). The optical fibre from the excitation module was then placed in the lens tube facing the power meter. The 520 nm and 638 nm lasers (Lasertack) were then recorded separately in continuous mode and under alternation (45 μs on, 55 μs off) for 15 hours and 1 hour, respectively.

#### Characterisation of the modulation timescales

The 520 nm and 638 nm lasers (Lasertack) on the setup were powered on and the alternation script from smOTTER was then run. A photodiode (Thorlabs) was connected to a lens tube (Thorlabs) and the fibre was placed in the tube. The voltage of the photodiode was recorded on an oscilloscope (Tektronix 2014B).

### 5.2. Accurate smFRET Measurements

Three duplex DNA standards were labelled with ATTO 550 23bp, 15bp and 11bp away from the acceptor dye, ATTO 647N (supplementary information). The DNA duplex samples were diluted to approximately 25 pM with an observation buffer (20 mM MgCl_2_, 5 mM NaCl, 5 mM Tris, pH 7.5). A volume of 10 μL was pipetted onto a coverslip enclosed in an airtight gasket and the data was recorded using the smfBox. Single-molecule analysis was performed using Anaconda with Jupyter Notebooks and using the FRETBursts python module. To account for the additional red laser power (0.6 mW) exceeding the commercial laser module power for common smFRET experiments, the alternation period for the red laser was reduced 15 μs / 100 μs cycles, whilst the green laser stayed on for 75 μs / 100 μs cycles. The background was corrected with a reduced threshold of L = 10 and F = 20, compared to previous studies to account for a lower laser power. The spectral cross-talk factors α and δ were determined by defining a donor only population >0.95 and an acceptor only population <0.2 (supplementary information). The stoichiometry and FRET efficiencies of the from all three DNA standards were plotted and fitted to obtain the last correction factors γ and β. The corrected FRET efficiencies were then determined using these four correction factors.

### 5.3. FCS measurements

The green laser was turned on and the Rhodamine 6G solution was diluted with water until the smfBox measured counts of ~25 kHz. A 2 minute acquisition was recorded. A known diffusion coefficient of 414 μm^2^/s for Rhodamine 6G was then used as a standard to determine the confocal volume. A dilute (~10 nM) doubly labelled DNA standard as previously described was placed on the smfBox using the same method and was recorded.

The diffusion coefficient and molecular brightness were then calculated in the MATLAB software package PAM.^[20]^ First, the signal from the observation volume was recorded over time and then correlated to provide an autocorrelation curve (Figure 6) using Equation 3.

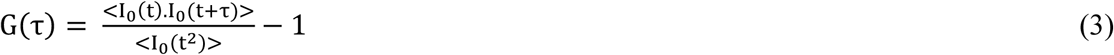

The decay of the autocorrelation curve depends on the rate of diffusion of the sample. A 3D diffusion model shown in Equation 4 was then used to fit this curve.

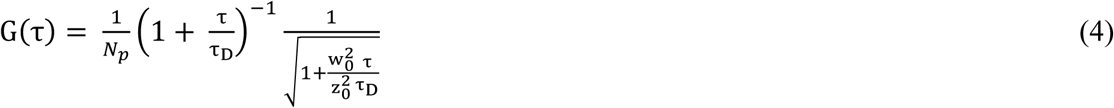

## Supporting Information

Supporting Information is available from the Wiley Online Library or from the author.

## Acknowledgements

This work was funded by: an EPSRC DTP studentship (TDC and AJC); a BBSRC Grant number (TDC), and an EPSRC Prize Fellowship (BA).

---

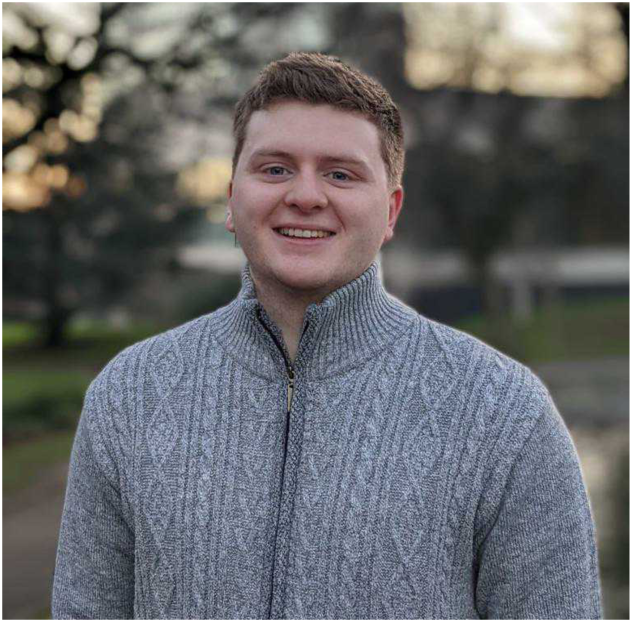

**Dylan George** is a Biophysics PhD student in the Craggs and Cadby labs at The University of Sheffield. He received his master’s degree in Chemistry in 2018 and has since been working towards designing single-molecule methods that are accessible through commercial and open-source projects. His current research focuses on developing a high-throughput single-molecule FRET instrument for drug discovery applications. His research interests include microscopy, single-molecule biophysics and imaging.

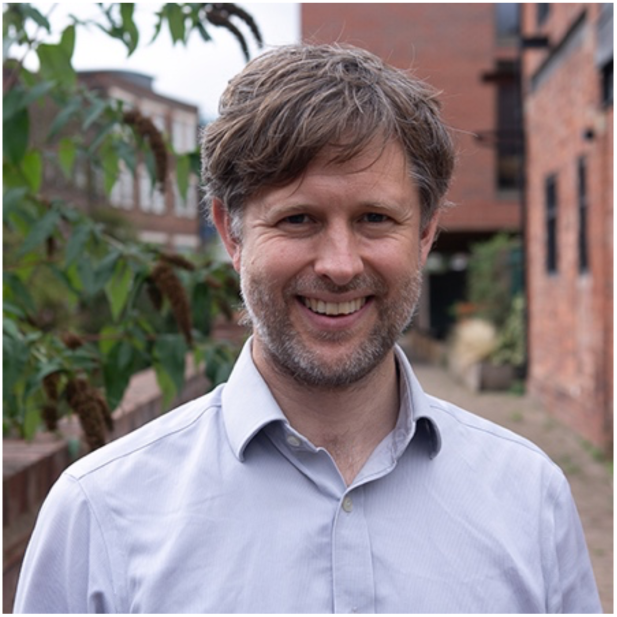

**Tim Craggs** runs an interdisciplinary lab in the University of Sheffield, focussing on the development and application of single-molecule fluorescence techniques to addressing crucial questions across physics, chemistry and the life sciences. He has pioneered the use of single-molecule Forster Resonance Energy Transfer (smFRET – a molecular ruler for the 30-90 Å scale) for the measurement of absolute distances with angstrom accuracy, opening the door to FRET driven structural biology. He is passionate about open science, and democratising powerful single-molecule methods, to make them available to the widest possible userbase, founding the spin-out company Exciting Instruments Ltd to realise this vision.

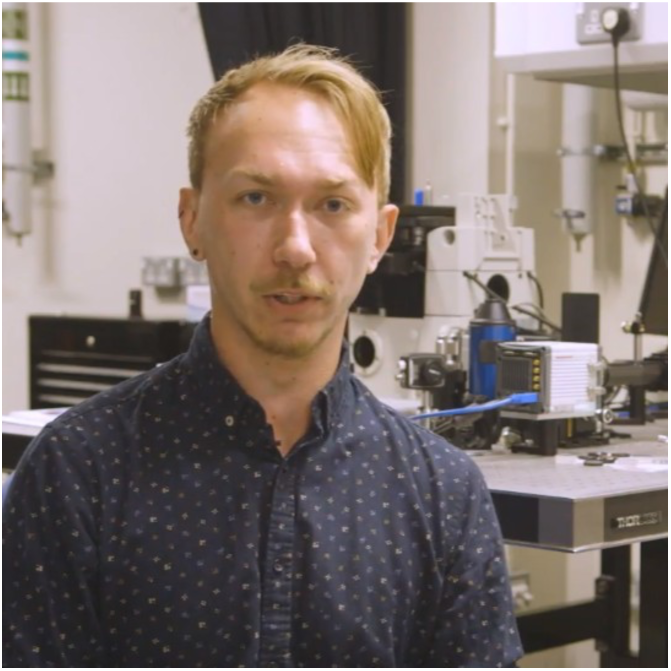

**Benjamin Ambrose** is an EPSRC Doctoral Prize Fellow working at the University of Sheffield, having completed his PhD in Biophysics in 2021. His research focuses on new tools for single-molecule fluorescence spectroscopy, such as developing open-source instrumentation for smFRET, and novel fluorescence quenching techniques for probing biomolecular interactions beyond the limits of typical FRET methods.

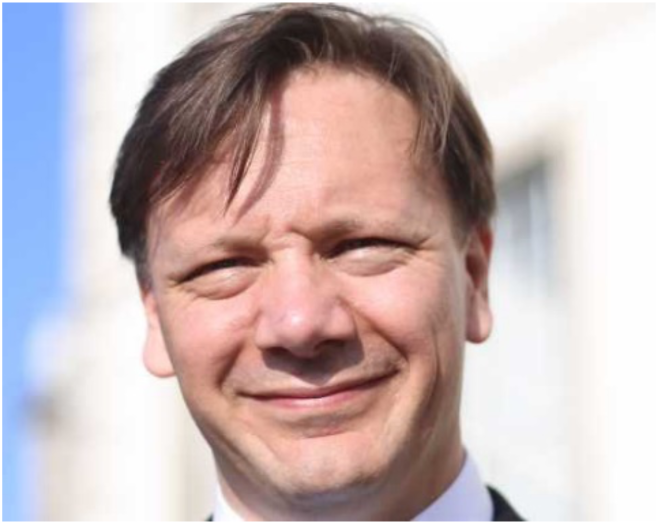

**Ashley Cadby** is a Professor of Biophysics in the Imagine Life consortium at the University of Sheffield. His research interests include Microscopy and Single molecule biophysics. The Cadby Lab sits at the core of the Bio Imaging centre, hosted within the Biophysics Research Group at the University of Sheffield, UK. The group is known for the development of optical systems for biological imaging focusing on their application to real world problems. This may be repurposing existing technology to build a better performing optical system, or the development of novel imaging to solve a specific problem.

## References

[1] G. E. Fantner and A. C. Oates, “Instruments of change for academic tool development,” Nature Physics, vol. 17, no. 4. pp. 421–424, 2021, doi: 10.1038/s41567-021-01221-3.

[2] R. Strack, “The miCube open microscope,” Nat. Methods, 2019, doi: 10.1038/s41592-019-0607-4.

[3] P. G. Pitrone et al., “OpenSPIM: An open-access light-sheet microscopy platform,” Nature Methods. 2013, doi: 10.1038/nmeth.2507.

[4] P. Almada et al., “Automating multimodal microscopy with NanoJ-Fluidics,” Nat. Commun., 2019, doi: 10.1038/s41467-019-09231-9.

[5] I. D. Schröder, J. Deschamps, A. Dasgupta, U. Matti, and J. Ries, “Cost-efficient open source laser engine for microscopy,” Biomed. Opt. Express, 2020, doi: 10.1364/boe.380815.

[6] P. R. Nicovich, J. Walsh, T. Böcking, and K. Gaus, “NicoLase - An open-source diode laser combiner, fiber launch, and sequencing controller for fluorescence microscopy,” PLoS One, 2017, doi: 10.1371/journal.pone.0173879.

[7] J. T. Collins et al., “Robotic microscopy for everyone: the OpenFlexure microscope,” Biomed. Opt. Express, 2020, doi: 10.1364/boe.385729.

[8] M. Ovesný, P. Křížek, J. Borkovec, Z. Švindrych, and G. M. Hagen, “ThunderSTORM: A comprehensive ImageJ plug-in for PALM and STORM data analysis and super-resolution imaging,” Bioinformatics, 2014, doi: 10.1093/bioinformatics/btu202.

[9] I. M. Müller, V. Mönkemöller, S. Hennig, W. Hübner, and T. Huser, “Open-source image reconstruction of super-resolution structured illumination microscopy data in ImageJ,” Nat. Commun., 2016, doi: 10.1038/ncomms10980.

[10] S. Culley et al., “Quantitative mapping and minimization of super-resolution optical imaging artifacts,” Nat. Methods, 2018, doi: 10.1038/nmeth.4605.

[11] B. Ambrose et al., “The smfBox is an open-source platform for single-molecule FRET,” Nat. Commun., vol. 11, no. 1, p. 5641, 2020, doi: 10.1038/s41467-020-19468-4.

[12] A. N. Kapanidis, T. A. Laurence, K. L. Nam, E. Margeat, X. Kong, and S. Weiss, “Alternating-laser excitation of single molecules,” Acc. Chem. Res., 2005, doi: 10.1021/ar0401348.

[13] O. Krichevsky and G. Bonnet, “Fluorescence correlation spectroscopy: The technique and its applications,” Reports Prog. Phys., vol. 65, no. 2, pp. 251–297, 2002, doi: 10.1088/0034-4885/65/2/203.

[14] P. Kask, P. Piksarv, and U. Mets, “Fluorescence correlation spectroscopy in the nanosecond time range: Photon antibunching in dye fluorescence,” Eur. Biophys. J., vol. 12, no. 3, pp. 163–166, 1985, doi: 10.1007/BF00254074.

[15] M. Bates, S. A. Jones, and X. Zhuang, “Stochastic optical reconstruction microscopy (STORM): A method for superresolution fluorescence imaging,” Cold Spring Harb. Protoc., vol. 8, no. 6, pp. 498–520, 2013, doi: 10.1101/pdb.top075143.

[16] N. K. Lee et al., “Accurate FRET measurements within single diffusing biomolecules using alternating-laser excitation,” Biophys. J., 2005, doi: 10.1529/biophysj.104.054114.

[17] J. Hohlbein, T. D. Craggs, and T. Cordes, “Alternating-laser excitation: Single-molecule FRET and beyond,” Chemical Society Reviews, vol. 43, no. 4. pp. 1156–1171, 2014, doi: 10.1039/c3cs60233h.

[18] B. Hellenkamp et al., “Precision and accuracy of single-molecule FRET measurements—a multi-laboratory benchmark study,” Nat. Methods, 2018, doi: 10.1038/s41592-018-0085-0.

[19] A. Ingargiola, E. Lerner, S. Y. Chung, S. Weiss, and X. Michalet, “FRETBursts: An open source toolkit for analysis of freely-diffusing Single-molecule FRET,” PLoS One, 2016, doi: 10.1371/journal.pone.0160716.

[20] W. Schrimpf, A. Barth, J. Hendrix, and D. C. Lamb, “PAM: A Framework for Integrated Analysis of Imaging, Single-Molecule, and Ensemble Fluorescence Data,” Biophys. J., 2018, doi: 10.1016/j.bpj.2018.02.035.

