## Supplementary material for "An Open-source, Cost-Efficient Excitation Module for Single-molecule Microscopy"

### 1. Supplementary Material

#### 1.1 Excitation module parts

| Part | Product information | Part | Product information |
|---|---|---|---|
| 2x Lasertack Laser Diode | 638 nm, (700 mW), 520 nm, (100 mW) | 45 deg Mirror Mount | H45A |
| 4x Table Clamp | CL5 | Dichroic Mirror | DMLP567T |
| 2x LED Heatsink (Fischer Elektronik) | SK 508 100 SA | FibrePort | PAF2A-A15A |
| 7x Universal Pole Holder | UPH1 | Post Mounting Bracket | HCP |
| 7x Optical Posts | TR30/M-P5 | Patch Cable, PM, 488 nm | P3-488PM-FC-2 |
| 6x Kinematic Mirror Mount | KM05/M | M6 Cap Screws | SH6MS16V |
| 5x Mirrors | PF05-03-G01 | Breadboard | MB3030/M |

**Table 1.** List of all the parts of the excitation module (all from Thorlabs unless stated otherwise).

#### 1.2 DNA Sequences

| Name | Sequence |
|---|---|
| 1a | 5' – GAG CTG AAA GTG TCG AGT TTG TTT GAG TGT TTG TCT GG – 3'<br>3' – CTC GAC TTT CAC AGC TCA AAC AAA CTC ACA AAC AGA CC – 5' – biotin |
| 1b | 5' – GAG CTG AAA GTG TCG AGT TTG TTT GAG TGT TTG TCT GG – 3'<br>3' – CTC GAC TTT CAC AGC TCA AAC AAA CTC ACA AAC AGA CC – 5' – biotin |
| 1c | 5' – GAG CTG AAA GTG TCG AGT TTG TTT GAG TGT TTG TCT GG – 3'<br>3' – CTC GAC TTT CAC AGC TCA AAC AAA CTC ACA AAC AGA CC – 5' – biotin |

**Table 2**. Accurate FRET standards used in the previous studies. Highlighted base pairs in red (acceptor) are labelled with ATTO 647N and the base pairs in green are labelled with ATTO 550. Biotin was also labelled for surface immobilisation for TIRF experiments, which were not used in this study.

#### 1.3 FRET Efficiency Data

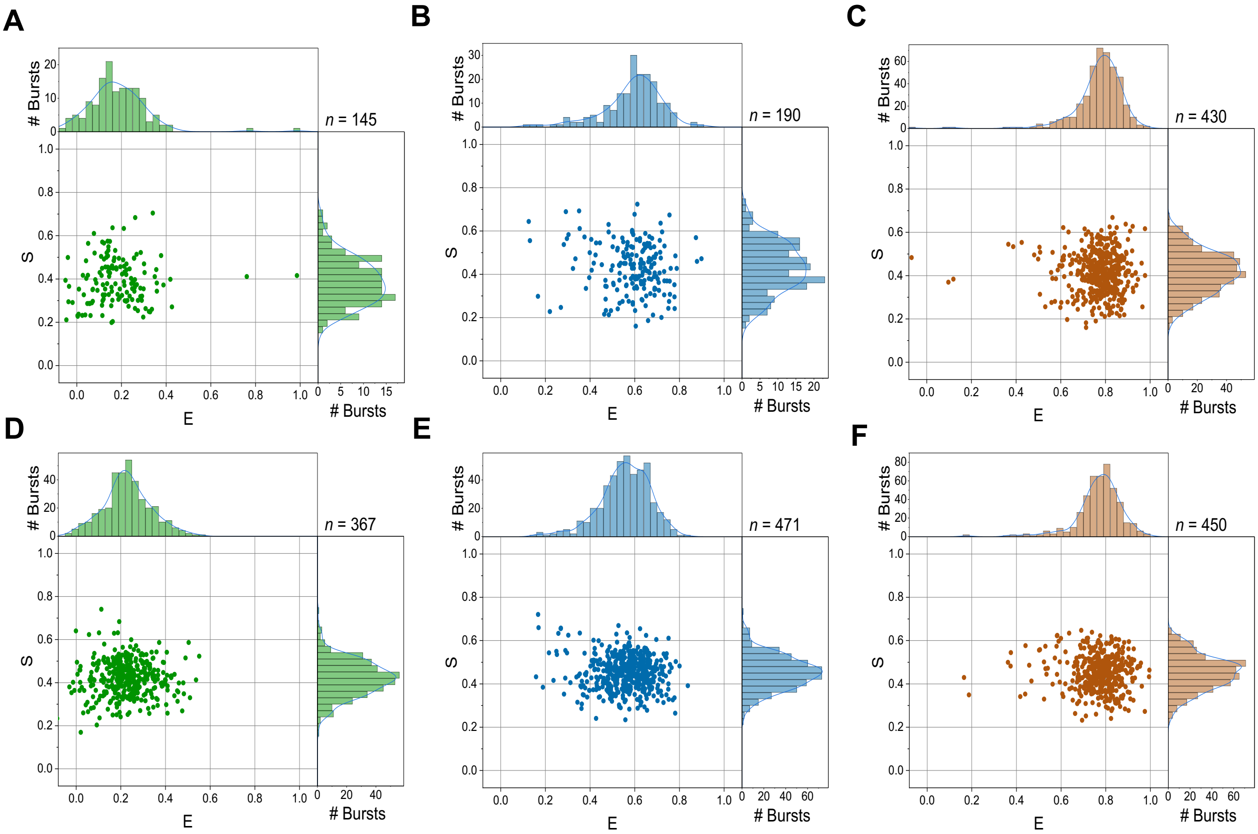


**Figure 1.** E-S histograms for the samples 1a, 1b and 1c. A-C) 1a-1c E-S histograms D-F) 1a-1c repeat experiment E-S histograms

**Table 1.** FRET Efficiencies of 1a, 1b and 1c

| **1a** | **1b** | **1c** |
|---|---|---|
| 0.225 | 0.570 | 0.784 |
| 0.171 | 0.628 | 0.793 |

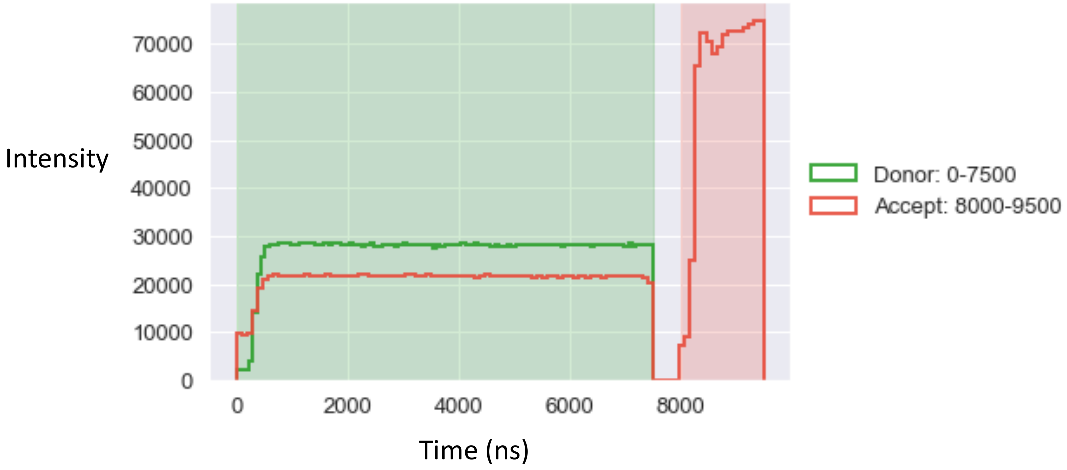


**Figure 2.** Alternation cycle of the laser diodes on the excitation module.

##### 1.4 Finding α and δ Correction factor

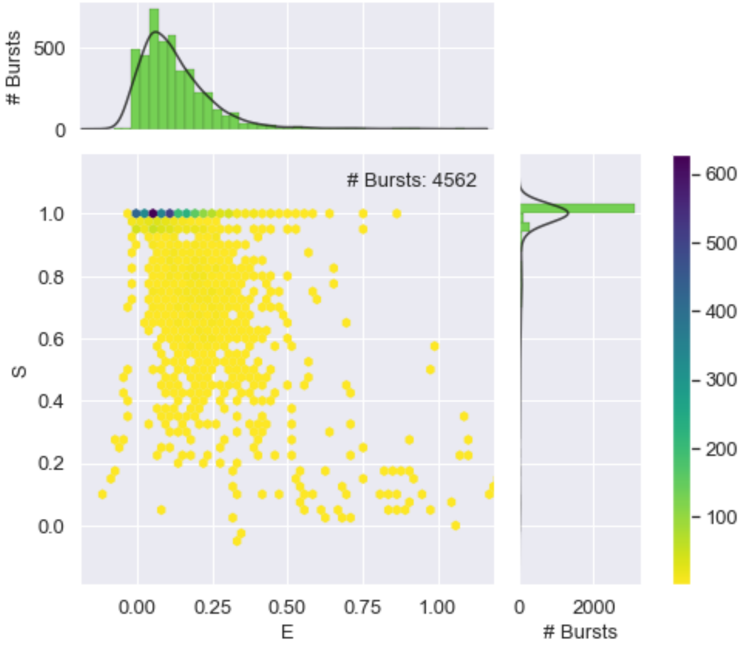


**Figure 3.** Uncorrected E-S histogram for sample 1a, used to determine bursts in the donor only and acceptor only region.

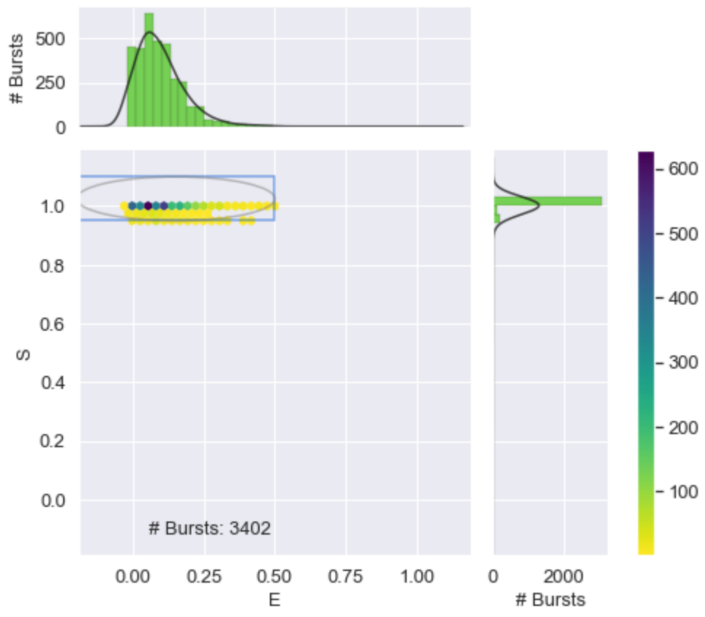


**Figure 4.** Uncorrected E-S histogram for sample 1a, highlighting the donor only region (S >0.95).

The donor only single-molecule events are used to calculate α (the photons emitted by donor detected by the acceptor detector) via the equation:

$$\alpha = \frac{E_{donor\ only}}{(1 - E_{donor\ only})} \tag{1}$$

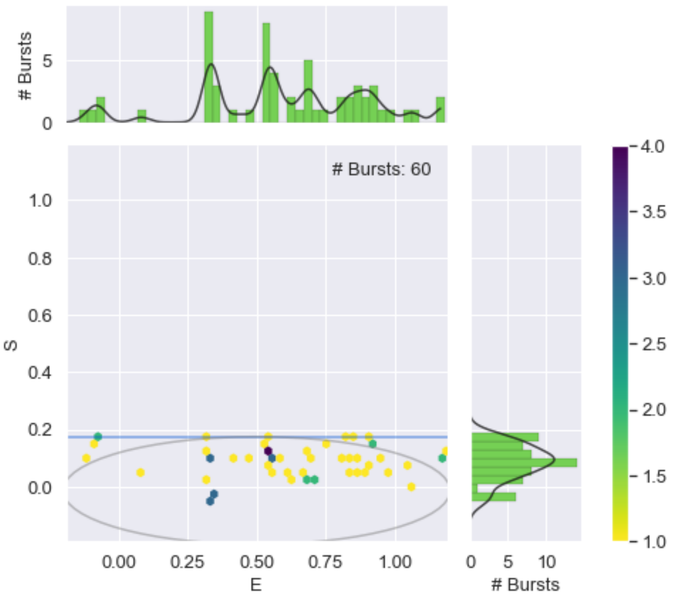

**Figure 5.** Uncorrected E-S histogram for sample 1a, showing the acceptor only region (S < 0.2).

The acceptor only single-molecule events are used to calculate δ (the direct excitation factor) via the equation:

$$\delta = \frac{S_{acceptor\ only}}{(1 - S_{acceptor\ only})} \quad (2)$$

##### 1.5 Finding γ + β correction factor

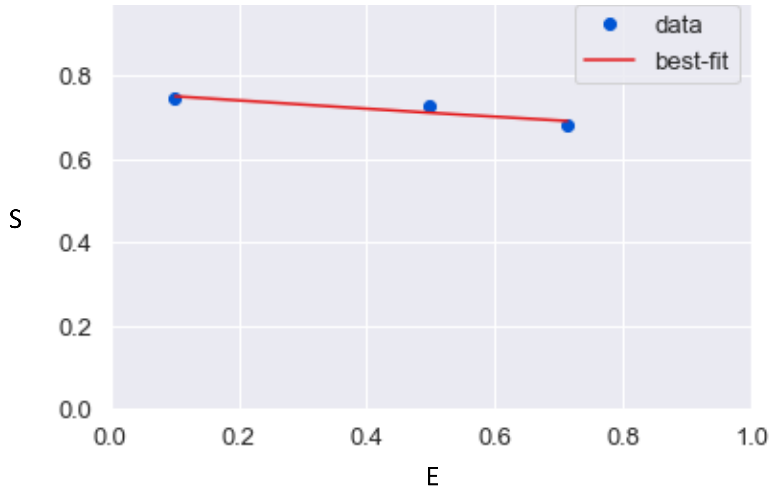


**Figure 6.** A graph showing the extracted E and S positions of all three duplex DNA standards.

Both γ and β correction factors can be determined by fitting the data to the following equation:

$$S = \frac{1}{1 + \beta\,\gamma + (1 - \gamma).\beta\,.E} \quad (3)$$

##### 1.6 FCS

**Table 2.** FCS data acquired from the MATLAB package PAM

| Sample | Counts [KHz] | Brightness [KHz] | N | D [$\mu m^2/s$] | $w_r$[µm] | $w_D$[µm] |
|---|---|---|---|---|---|---|
| **Rhodamine 6G** | 58.2 | 5.94 | 9.80 | 118.6 | 0.73 | 2.11 |
| **Duplex DNA (1c)** | 5.04 | 2.32 | 2.17 | 414 | 0.73 | 2.11 |
